# A Cre-responsive reporter quail for flexible genetic lineage tracing

**DOI:** 10.64898/2026.09.20.751349

**Authors:** Siew Zhuan Tan, Hila Barzilai-Tutsch, Haolan Sun, Ye-Wheen Lim, Christophe Marcelle, Olivier Serralbo, Melanie D. White

## Abstract

Genetic lineage tracing is a powerful approach for investigating cell fate and tissue development, but versatile Cre/loxP-based tools remain limited in avian models. Here, we generated and characterised a Cre-responsive transgenic quail line that enables permanent genetic labelling of embryonic cell populations. The reporter line ubiquitously expresses nuclear-localised mCherry, which is excised following Cre-mediated recombination, enabling expression of nuclear-localised EGFP. We demonstrate that this reporter can be coupled to multiple Cre delivery strategies, providing flexible spatial and temporal control of lineage labelling. Ubiquitous or cell-type-specific Cre expression induces recombination and stable EGFP labelling that is maintained during clonal expansion. Optogenetic activation of a split Cre system enables highly spatially and temporally restricted recombination. Furthermore, lipid nanoparticle-mediated delivery of Cre mRNA enables efficient recombination in distinct embryonic cell populations, including mesodermal, neuroepithelial, neural crest, endothelial and circulating cells. This transgenic reporter quail provides a flexible platform for permanent lineage labelling without the need to generate tissue-specific Cre transgenic lines and can be readily combined with diverse Cre delivery strategies to investigate cell fate, lineage and morphogenesis *in vivo*.

**SUMMARY STATEMENT:** This new transgenic quail line provides a flexible way to permanently label cells during embryo development, enabling researchers to follow their fate and behaviour over time.

## INTRODUCTION

Lineage tracing is a fundamental approach in developmental biology for understanding cell fate by labelling a defined progenitor population and subsequently tracking its descendants over time. Avian embryos have played an important role in establishing developmental fate maps, particularly through the use of quail-chick chimeras to trace the contributions and migration of embryonic cell populations (Le Douarin, 1973). This cell transplantation approach has provided fundamental insights into diverse developmental processes, including neural crest migration, tissue patterning and organ development (Ribatti, 2019). Several approaches have subsequently been developed for lineage labelling in avian embryos; however, each has limitations for long-term lineage tracing. Vital dyes such as DiI can label cells for several hours to days of development (Chinnaiya et al., 2023, Jacobs-Li et al., 2023), but the label becomes progressively diluted through cell division. Retroviral reporter vectors provide more stable labelling through genomic integration, enabling the label to be inherited by descendant cells (Gandhi et al., 2021). However, this approach requires approximately two days for robust reporter protein expression to become detectable and depends on sustained reporter expression following viral integration. The Cre/loxP system provides an alternative strategy in which expression of Cre recombinase induces irreversible recombination of a reporter, thereby permanently marking the progenitor population and its descendants. This approach is widely used for lineage tracing in mouse and zebrafish models (Ming et al., 2026, Takahashi et al., 2024) and has more recently been demonstrated in the Chameleon transgenic chicken line (Oh et al., 2024). However, a comparable Cre-responsive transgenic reporter line has not yet been established in quail.

Japanese quail (*Coturnix japonica*) offers several practical advantages as an avian model for developmental studies. Like chicken (*Gallus gallus*), quail embryos develop within externally laid eggs, making them readily accessible for experimental manipulation and real-time observation of dynamic developmental processes. Quail are also smaller in size and reach sexual maturity in approximately six weeks, compared with approximately six months for chicken, resulting in a shorter generation time and reducing the cost of maintaining and expanding transgenic lines. Despite these differences, quail and chicken show high conservation in genomic organisation, including chromosomal synteny, gene content and gene orthologs (Morris et al., 2020). The development of transgenic quail lines has further expanded the experimental capabilities of this model, enabling studies of cell dynamics, tissue mechanics and morphogenesis (Alvarez et al., 2024, Barzilai-Tutsch et al., 2022, Caldarelli et al., 2024, Harn et al., 2026, Huss et al., 2015, Serralbo et al., 2020, Villedieu et al., 2025, Wang et al., 2026). Together, these studies highlight the potential of transgenic quail as a versatile platform for developmental studies and provide a strong foundation for extending quail transgenesis to permanent lineage tracing.

Here, we generated a Cre-responsive reporter quail line, TgT2[CAG:lox-nlsmCherry-lox-nlsEGFP], which enables permanent genetic labelling of embryonic cell populations. The line ubiquitously expresses nuclear-localised mCherry, which is excised upon Cre-mediated recombination, allowing the expression of nuclear-localised EGFP. We first validated the floxed reporter line by delivering a ubiquitously expressed Cre vector through electroporation. We then exploited a cell type-specific enhancer to induce Cre-mediated recombination in neural crest cells. We further characterised the transgenic line using optogenetic activation of Cre recombinase and lipid nanoparticle (LNP)-mediated delivery of Cre mRNA. Together, our results establish a versatile Cre-responsive reporter line that can be combined with multiple Cre recombinase delivery strategies to enable spatially and temporally flexible genetic lineage labelling in avian embryos.

## RESULTS AND DISCUSSION

### Generation of a CAG:lox-nlsmCherry-lox-nlsEGFP transgenic quail line

We generated a transgenic quail line expressing a Cre-responsive reporter under the control of the strong ubiquitous promoter (CAG: CMV enhancer and chicken β-actin promoter) (Hitoshi et al., 1991). The CAG:lox-nlsmCherry-lox-nlsEGFP cassette was flanked by 5’ and 3’ Tol2 (T2) transposable elements to enable stable integration into the quail genome (Fig. 1A). A transfection mixture containing lipofectamine 2000, the CAG:lox-nlsmCherry-lox-nlsEGFP vector and a CAG:Transposase construct was injected into the dorsal aorta of Hamilton-Hamburger stage (HH) 15-16 (E2-2.5) quail embryos to transfect circulating primordial germ cells (Alvarez et al., 2024, Barzilai-Tutsch et al., 2022, Serralbo et al., 2020, Tyack et al., 2013). The founders were identified and mated with wild-type quails to establish stable lines (TgT2[CAG:lox-nlsmCherry-lox-nlsEGFP]). They are viable, phenotypically normal, and fertile and can be maintained as either homozygotes or heterozygotes. After further breeding the lines were indistinguishable and a single line was selected for long-term maintenance. Robust nuclear mCherry expression was detected from stage HH5 (head-process, HP) onwards (Hamburger and Hamilton, 1951), with Phalloidin staining outlining the embryo morphology (Fig. 1B). Nuclear mCherry expression remained stable even after five generations of breeding.

**Fig 1.**
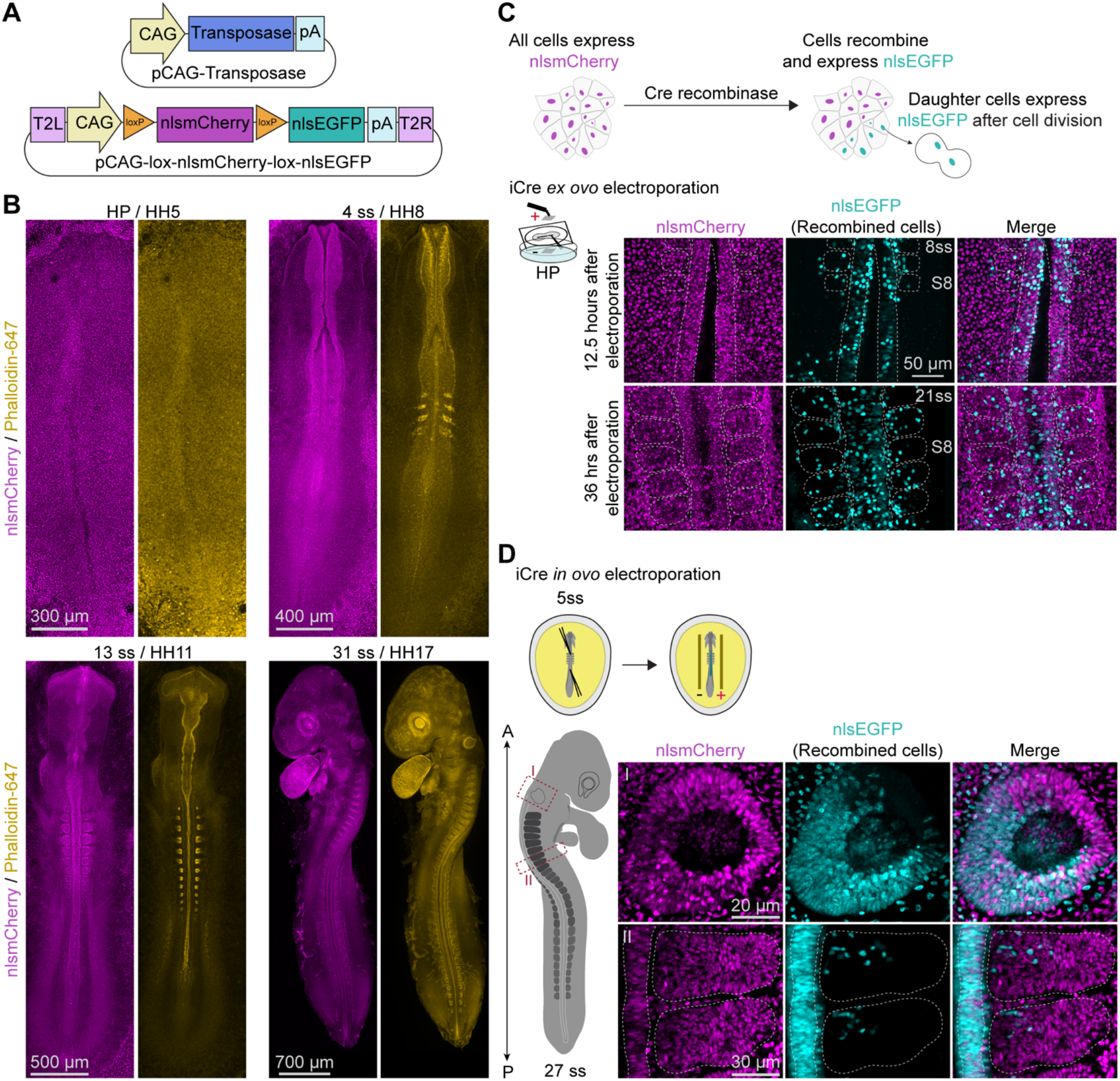
Generation and characterisation of TgT2[CAG:lox-nlsmCherry-lox-nlsEGFP] transgenic quail embryo. (A) Plasmid vectors were used to generate the transgenic line. (B) Whole-mount confocal images of transgenic embryo at the indicated developmental stages. Phalloidin-647 staining (gold) outlines embryonic morphology, and nuclear mCherry is shown in magenta. (C) A schematic diagram showing Cre recombination mechanism. Electroporated cells with iCre plasmid recombine after 12.5 hours (top panel). Stable nuclear EGFP expression and clonal expansion of recombined cells at the same somite level (S8) following cell division (bottom panel). (D) Cre-mediated recombination is detected in the otic vesicle and neural tube following *in ovo* electroporation of iCre plasmid.

To deliver the Cre recombinase, embryos were electroporated *ex ovo* at HP with a plasmid expressing *iCre* under the control of the CMV promoter. The plasmid was injected into the space between the neural plate and vitelline membrane, caudal to Hensen’s node, and cells within this region were targeted by electroporation. Cre-mediated recombination, indicated by activation of nuclear EGFP expression, was detected 12.5 hours after electroporation (Fig. 1C, top panel). Stable nuclear EGFP expression was maintained during clonal expansion (Fig. 1C, bottom panel), validating the reporter for lineage tracing applications (Loulier et al., 2014, Oh et al., 2024, Schick et al., 2019). Cre-mediated recombination was also detected in multiple tissues, with nuclear EGFP-positive cells observed in the neuroepithelium, otic vesicle (Fig. 1D), and dorsal root ganglion (DRG) (Fig. S1).

To validate the specificity of Cre-mediated recombination, we performed *ex ovo* electroporation using HP embryos with either both iCre and H2B-TagBFP2 constructs or the H2B-TagBFP2 construct alone (Fig. S2A). The embryos were fixed at 24 hours after electroporation and stained for Cre. Embryos electroporated with only the H2B-TagBFP2 plasmid showed neither recombined nuclear EGFP-positive cells nor Cre-positive cells by immunofluorescence (Fig. S2B, bottom panel), indicating that recombination does not occur in the absence of Cre recombinase. Although not all recombined nuclear EGFP-positive cells retained detectable Cre expression 24 hours after electroporation (Fig. S2B, top panel), this result suggests that a transient, and potentially low level of Cre recombinase is sufficient to induce permanent recombination. We also observed residual nuclear mCherry expression in the recombined cells despite robust nuclear EGFP expression. This likely reflects the persistence of pre-existing mCherry protein following Cre-mediated excision, consistent with the intrinsic stability of mCherry (Shaner et al., 2004) and a phenomenon commonly observed with fluorescent protein-based Cre reporters across different organisms (Hartwich et al., 2012, Yoshinari et al., 2012).

### Cre-mediated recombination in neural crest cells

Neural crest cells are a transient, multipotent vertebrate cell population that arises at the neural plate border and undergoes epithelial-to-mesenchymal transition (EMT) followed by extensive migration (Graham, 2003). They can be subdivided into cranial, vagal, trunk, and sacral populations according to their axial origins, and give rise to diverse derivatives, including craniofacial cartilage, parasympathetic ganglia, melanocytes, dorsal root ganglia, and pelvic ganglia. Their migration and differentiation are regulated by a network of transcription factors, such as Snail/Slug, FoxD3, and Sox10 (Betancur et al., 2010a, Williams et al., 2019). Most studies investigating neural crest development rely on transient enhancer-driven reporters (Betancur et al., 2010b, Gandhi et al., 2026, Murko and Bronner, 2017), in which reporter expression is diluted over time due to cell division. Therefore, the generation of a Cre-responsive avian reporter line provides a powerful approach for permanent labelling.

Here, we used a previously characterised vagal/trunk neural crest cell-specific enhancer, *FoxD3* NC2 enhancer (Simões-Costa et al., 2012), to drive Cre-mediated recombination. The enhancer was cloned upstream of *iCre* under the control of the minimal TK promoter (Fig. 2A). Embryos were electroporated unilaterally *in ovo* at the 6-somite stage (ss) (Fig. 2B), resulting in plasmid uptake and Cre-mediated recombination predominantly on the right side of the embryo (Fig. S3). NC2 enhancer-driven recombination was primarily observed in the migratory trunk neural crest cells (Fig. 2C), as confirmed by Sox10 immunostaining. Interestingly, we observed heterogenous nuclear EGFP expression among NC2 enhancer-induced recombined cells, with some cells exhibiting stronger and others weaker fluorescence, whereas CMV promoter-driven *iCre* resulted in relatively uniform nuclear EGFP expression (Fig. 1D and Fig. S1). This difference suggests that the heterogeneity in reporter expression may be associated with differences in NC2 enhancer activity rather than variability in the Cre/loxP recombination system itself. Given that FoxD3 is a developmentally-regulated transcription factor and NC2 functions as a developmental enhancer, this heterogeneity may reflect the dynamic regulation of NC2 enhancer activity during neural crest specification. Such heterogeneity is consistent with previous findings (Azambuja and Simões-Costa, 2021, Betancur et al., 2010b, Murko and Bronner, 2017) that neural crest cells comprise transcriptionally distinct subpopulations and that enhancer activity can encode spatial and temporal information associated with neural crest specification.

**Fig 2.**
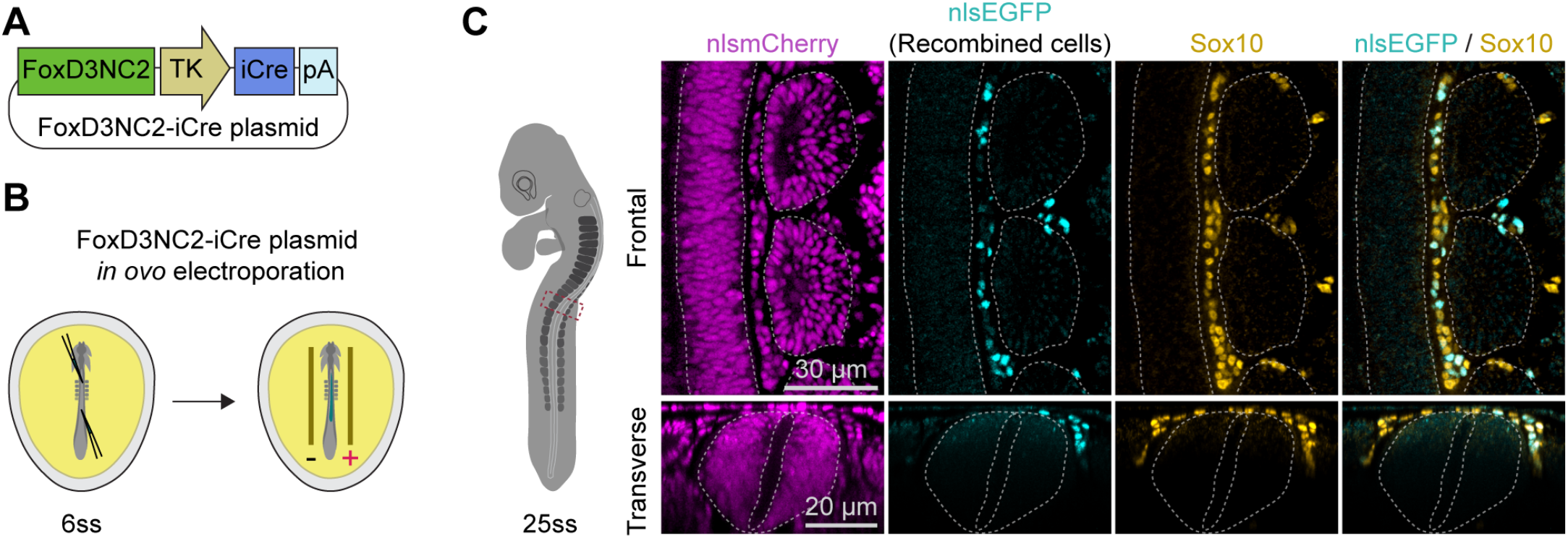
Cre-mediated recombination induced by the *FoxD3* NC2 enhancer. (A) Plasmid vector used for *in ovo* electroporation. (B) A schematic diagram showing unilateral *in ovo* electroporation perform at 6ss, with the right side electroporated. (C) Frontal and transverse views of a 25ss embryo showing recombined nuclear EGFP-positive cells driven by NC2 enhancer. Recombined cells exhibit ventrolateral migration (transverse view) and co-localise with Sox10-positive neural crest cells (gold).

A previous study used another neural crest cell-specific enhancer, *FoxD3* #168, identified from the Vista enhancer browser, to drive Cre expression in embryos electroporated with a CAG:lox-STOP-lox-EGFP reporter construct (Nitzan et al., 2013). Other studies have also applied cell type-specific enhancers, including enhancers active in hindbrain interneurons (Kohl et al., 2013) and spinal interneurons (Hadas et al., 2014), to induce Cre-mediated recombination in avian embryos. These lineage labelling approaches primarily relied on co-electroporation of Cre-expressing and loxP-based reporter plasmids or PiggyBac-mediated genomic integration. In contrast, our approach utilises a germline transgenic floxed reporter, eliminating the requirement for co-electroporation of reporter constructs and providing a stable, genetically encoded platform for lineage labelling. Furthermore, the presence of nuclear mCherry reporter in the same genetic background enables simultaneous visualisation of tissue morphology and recombined nuclear EGFP-positive cells during live imaging. To our knowledge, this is the first reported use of a cell type-specific enhancer to drive Cre-mediated recombination in a germline transgenic quail model, providing a valuable platform for permanent labelling and long-term lineage tracing of neural crest derivatives.

### Optogenetic induction of Cre-mediated recombination

Electroporation of the CMV-iCre construct established that our reporter is Cre-responsive, while the *FoxD3* NC2-iCre construct demonstrated that the system can be used for cell type-specific lineage labelling. However, neither approach provides experimentally controlled temporal regulation of Cre activity and therefore offers limited control over the onset of recombination. To achieve precise spatiotemporal control of lineage labelling, we next investigated whether our reporter line could be coupled with a light-inducible Cre recombinase (opto-Cre). We employed an opto-Cre system in which the N- and C-terminal fragments of Cre (CreN and CreC) were fused to the light-responsive proteins cryptochrome 2 (CRY2) and CRY2-interacting basic-helix-loop-helix protein 1 (CIB1), respectively, with the two fusion proteins expressed from a bicistronic construct separated by an IRES (Fig. 3A). Blue-light illumination induces CRY2-CIB1 dimerization, bringing CreN and CreC into proximity to reconstitute functional Cre recombinase and thereby induce recombination.

**Fig 3.**
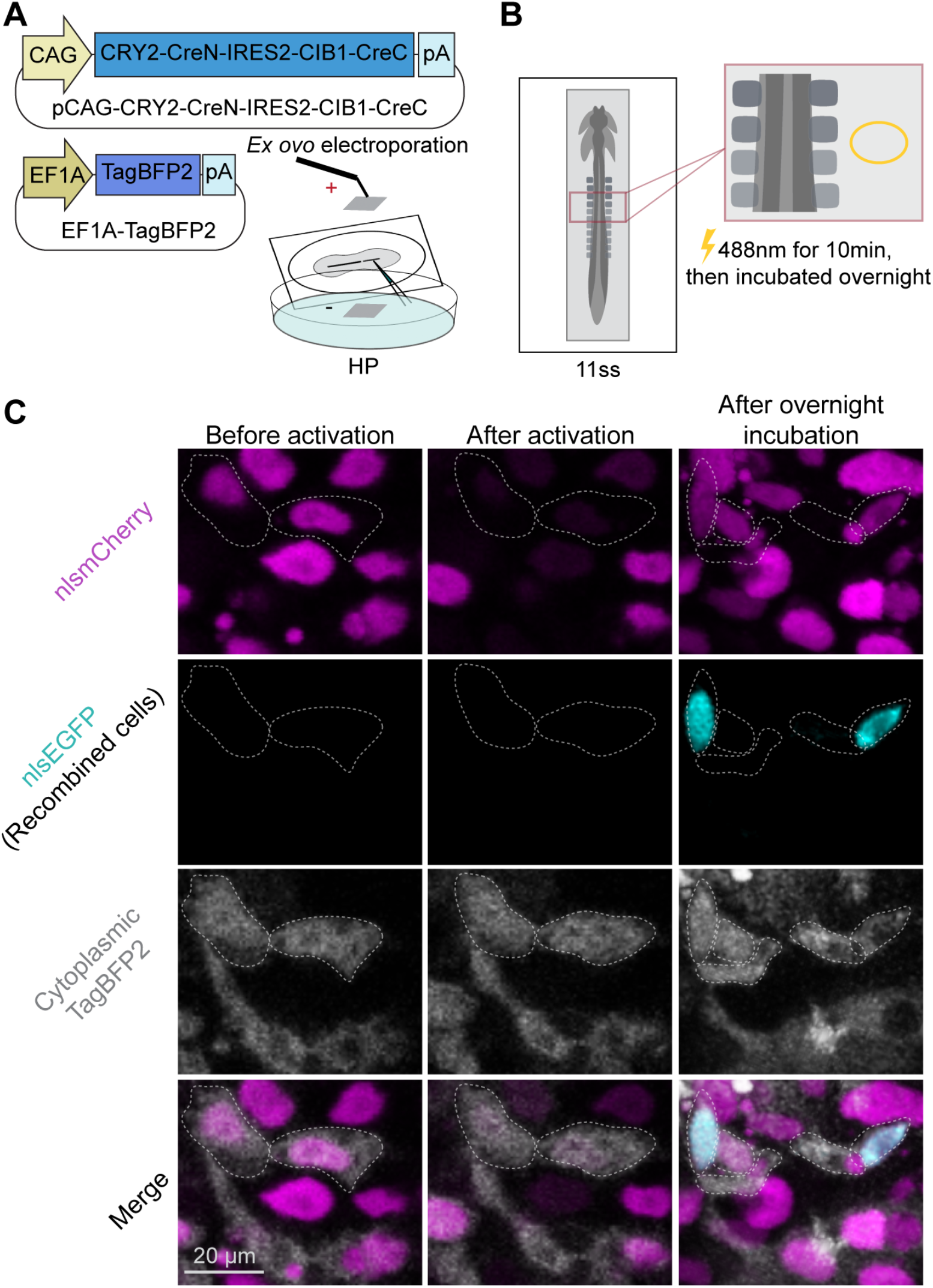
Spatially and temporally controlled Cre recombination by optogenetic activation. (A) Embryo is electroporated with the plasmid vectors through *ex ovo* electroporation. (B) A schematic diagram showing after overnight incubation, a region of interest (yellow circle) within the epidermis of the electroporated embryo is selected and exposed to 488 nm laser for 10 min. (C) Representative images showing before photoactivation, immediately after activation, and following overnight incubation.

To test whether this system could induce recombination in our reporter line, embryos were co-electroporated *ex ovo* at HP with the opto-Cre construct and a cytoplasmic TagBFP2 vector to visualise the electroporated area. Following overnight incubation, a region of the epidermis was selected and illuminated with 488 nm laser for ten minutes (Fig. 3B), after which the embryos were returned to the incubator for a further overnight incubation in dark conditions. Throughout the experiment, embryos were kept in the dark except during photoactivation to minimise potential self-activation of the opto-Cre system by ambient light. Before photoactivation, no recombined nuclear EGFP-positive cells were detected in the selected region (Fig. 3C). Immediately following photoactivation, nuclear mCherry fluorescence within the illuminated region was markedly reduced, consistent with photobleaching caused by the 488 nm illumination. Following overnight incubation, recombined nuclear EGFP-positive cells were detected within the illuminated region, demonstrating that photoactivation of opto-Cre was sufficient to induce Cre-mediated recombination. Residual nuclear mCherry expression was also observed in these cells, consistent with persistence of pre-existing mCherry protein following recombination.

The feasibility of light-inducible Cre-mediated gene activation in avian embryos has also recently been demonstrated using an alternative optogenetic system (Pfann et al., 2025). In this study, the Magnet-Cre system was applied in chicken embryos and demonstrated light-dependent gene activation following electroporation into the neural tube, including spatially restricted activation using localized blue-light illumination. This provides an independent demonstration that optogenetic Cre systems can be used to achieve spatial and temporal control of gene activation in developing avian embryos. Our CRY2/CIB1 system similarly enabled localized light-induced recombination in the transgenic quail reporter line but differs in its light-responsive components and reporter configuration. Together, these studies support the feasibility of optogenetic control of Cre-dependent gene activation in developing avian embryos.

Although photoactivation resulted in a lower frequency of recombination than that observed with constitutively expressed CMV-iCre and *FoxD3* NC2-iCre, recombination was readily detectable and, importantly, could be spatially restricted through targeted illumination. The efficiency of this approach is likely influenced by the mosaic nature of plasmid electroporation, the requirement for light-dependent association of the split Cre fragments, and the intracellular abundance and localisation of the individual components. While further optimisation of construct design, nuclear localisation and light-delivery parameters may increase recombination efficiency, the primary strength of this approach lies in achieving exceptionally precise spatial and temporal control over recombination.

This level of control is particularly valuable for lineage-tracing applications in which the objective is to label individual cells or very small groups of cells rather than an entire tissue. Targeted photoactivation provides the opportunity to mark cells at defined positions and developmental stages and subsequently follow their descendants, enabling lineage relationships and clonal behaviour to be resolved at a level of spatial precision that is difficult to achieve with conventional Cre-based approaches.

### LNP-mediated delivery of Cre mRNA enables robust reporter recombination

Having established that our reporter undergoes Cre-mediated recombination following constitutive, cell type-specific and light-inducible Cre recombinase expression, we next investigated whether recombination could be achieved through transient delivery of Cre messenger RNA (mRNA). Lipid nanoparticles (LNPs) provide an efficient platform for intracellular mRNA delivery by protecting RNA from degradation, facilitating cellular uptake, promoting endosomal escape, and enabling cytoplasmic release. We therefore tested whether LNP-mediated delivery of Cre mRNA could induce recombination in our reporter line.

We injected Cre mRNA LNPs (Fig. 4A) at different developmental stages and anatomical sites to target distinct cell populations. To label mesodermal cells, LNPs were injected into the posterior primitive streak. After overnight incubation, embryos were fixed and stained for Fibronectin, an extracellular matrix protein that surrounds the somites, which are derived from mesoderm or Brachyury, a mesodermal marker. LNP-mediated Cre delivery resulted in extensive recombination of the mesodermal tissues (Fig. 4B). To target neuroepithelial and neural crest cells, LNPs were injected into the neural tube lumen at 5ss (Fig. 4C). Cre mRNA LNP delivery resulted in recombination in the DRG and sympathetic ganglion (SG) (Fig. 4C, transverse view), as well as in the neuroepithelium (Fig. S4). The identity of recombined nuclear EGFP-positive cells was confirmed by staining for either Sox10, a neural crest marker, or Sox2, a marker of neural progenitor cells. Furthermore, injection of Cre mRNA LNPs into the yolk sac vasculature at day 3 of incubation resulted in extensive recombination in endothelial cells and circulating cells after overnight incubation (Fig. S5). Taken together, these results demonstrate that delivery of Cre mRNA via LNPs can induce recombination across multiple embryonic cell populations and can be adapted to target distinct tissues by varying the injection site and developmental stage. This is consistent with a previous study showing that LNP-mediated delivery of Cre mRNA can generate functional Cre recombinase and induce loxP-mediated genomic recombination in the mouse embryo (Tombácz et al., 2021). Importantly, this approach also avoids the need for electroporation, providing a less invasive strategy for delivering Cre recombinase to the developing embryo.

**Fig 4.**
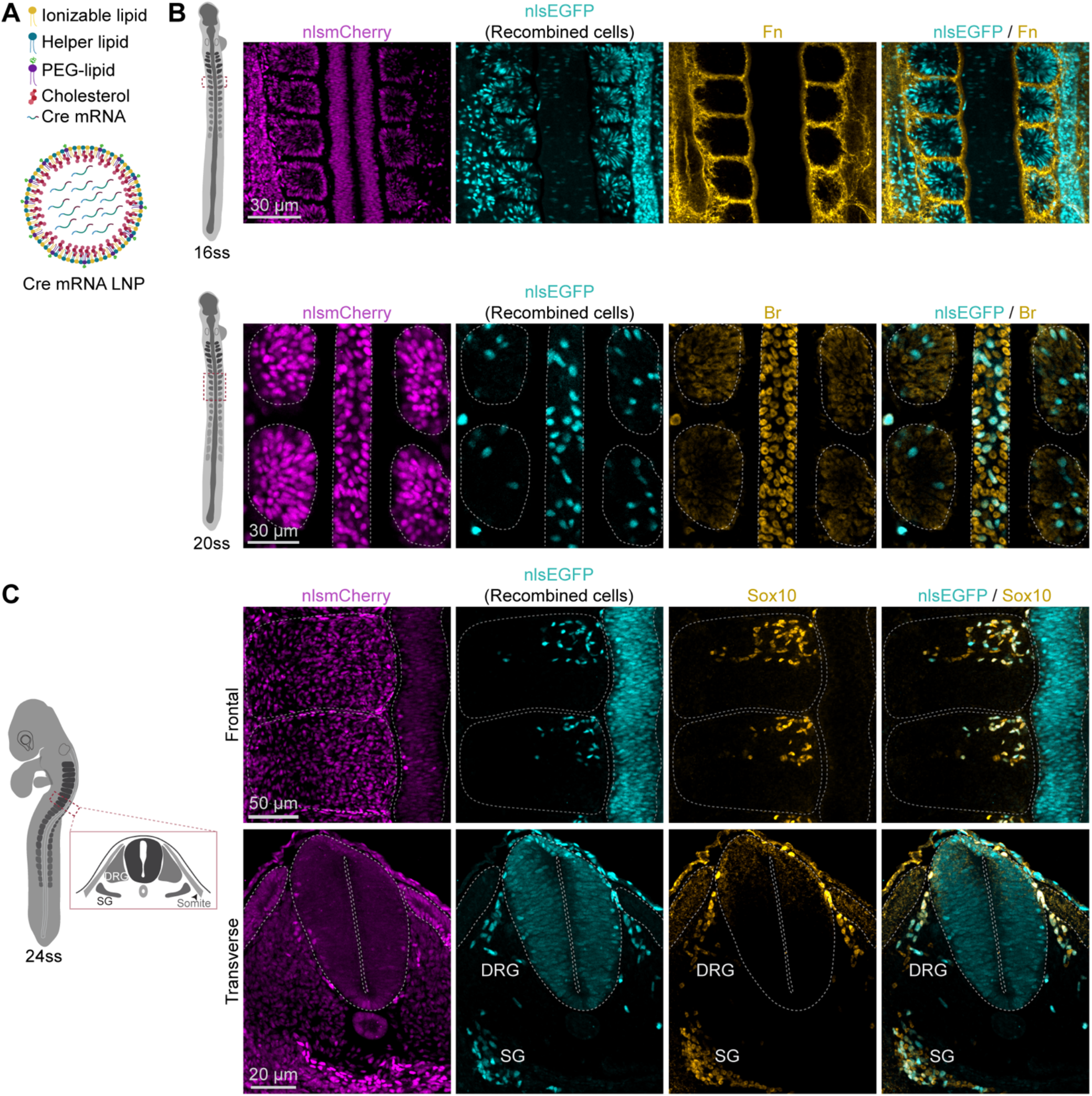
Cre mRNA LNP-mediated recombination. (A) A schematic diagram showing the composition of Cre mRNA LNPs. (B) Embryos were injected with Cre mRNA LNPs at PS and fixed after overnight incubation. Representative embryos at 16 and 20ss are shown. PS injection favoured recombination in mesodermal cells, with the identity and localisation of recombined nuclear EGFP-positive cells assessed by immunostaining for Fibronectin (Fn), which outlines the somite structures and Brachyury (Br), a mesodermal marker. (C) Embryos were injected with Cre mRNA LNPs at 5ss and fixed after overnight incubation. A representative embryo at 24ss is shown. Recombined nuclear EGFP-positive cells were detected in Sox10-expressing populations, including cells within the dorsal root ganglia (DRG) and sympathetic ganglia (SG).

Another transient approach for inducing Cre-mediated recombination in avian embryos was demonstrated by Oh et al. (2024), who used cell-permeable TAT-Cre protein delivered via Affi-Gel Blue beads in the Chameleon transgenic chicken line to induce recombination in discrete regions of the limb bud. Whereas TAT-Cre beads provide highly localized and transient Cre activity, our LNP-mediated Cre mRNA delivery provides a complementary strategy for transient Cre activity, in which different embryonic cell populations can be targeted by varying the route, anatomical site, and developmental timing of administration.

### CONCLUSIONS

Collectively, our results establish the TgT2[CAG:lox-nlsmCherry-lox-nlsEGFP] floxed reporter quail as a versatile platform for lineage labelling and conditional genetic manipulation in the avian embryo. Recombination can be achieved through a range of complementary approaches, from conventional electroporation of ubiquitously expressed or tissue-specific Cre to optogenetic activation of electroporated split-Cre, providing precise spatial and temporal control over recombination. LNP-mediated delivery of Cre mRNA further enables widespread recombination across multiple tissue types, with the distribution of labelled cells determined by the timing and site of delivery. Importantly, this flexibility enables lineage labelling without the need to generate or cross with tissue-specific Cre transgenic lines, considerably simplifying the application of genetic approaches in the quail embryo. Coupled with the accessibility of avian embryos to manipulation and live imaging, this reporter line provides a powerful and broadly applicable resource for following cell fate and lineage while simultaneously resolving the dynamic cellular behaviours that drive tissue morphogenesis *in vivo*.

## MATERIALS AND METHODS

### Generation of transgenic CAG:lox-nlsmCherry-lox-nlsEGFP quail line

The direct injection technique was performed as described previously (Alvarez et al., 2024, Barzilai-Tutsch et al., 2022, Serralbo et al., 2020, Tyack et al., 2013). The injection mix contained 0.6 µg of pCAG-lox-nlsmCherry-lox-nlsEGFP Tol2 plasmid, 1.2 µg of CAG Transposase plasmid, and 3 µL of Lipofectamine 2000 CD (Thermo Fisher Scientific) in 90 µL of OptiPro. About 1 µL of injection mix was injected in the dorsal aorta of 2.5-day-old embryos. Eggs were then sealed and incubated until hatching. Chicks were grown for six weeks until they reached sexual maturity. Semen from the G0 males was collected using a female teaser and massage techniques as described previously (Serralbo et al., 2020). Genomic DNA from semen was extracted and PCR genotype was performed to test for the presence of the transgene. Males showing a positive band were kept and crossed with wild-type females. Founders G1 offspring were selected using RFP goggles for the expression of ubiquitously expressed mCherry visible in chicks at hatching point.

### Maintenance of quails

Transgenic quails were housed and bred at the University of Queensland according to local animal ethical policies approved by the University of Queensland Health Science Animal Ethics Committee (2023/AE000559). Fertilised quail eggs were collected daily by the animal facility and kept at 14°C before use.

### Plasmid constructs

The CMV-iCre vector used in the experiment can be obtained from Addgene (Plasmid #112683). A *FoxD3* NC2-EGFP vector was generously provided by Bronner lab (Simões-Costa et al., 2012). A synthetic DNA fragment containing the required partial TK promoter and chimeric intron sequences together with the iCre coding sequences was synthesized by Twist Bioscience and supplied in the pTwist-Amp vector. Both vector constructs were digested with *MluI* and *XbaI*. The desired synthetic fragment was ligated into the *MluI* and *XbaI* sites of the *FoxD3* NC2 vector backbone to generate the *FoxD3* NC2-iCre vector. The final vector was verified by *BglII* restriction digestion and confirmed by Sanger sequencing. The opto-Cre vector was generated using Gibson assembly. An IRES2 fragment was added downstream of CRY2-CreN (Addgene, Plasmid #75368 using the primers listed in Table S1. The resulting CRY2-CreN-IRES2 fragment was then inserted into a vector backbone containing the CAG promoter using *SacI* and *MfeI*.CIBN-CreC (Addgene, Plasmid #75367 was subsequently inserted downstream of CAG-CRY2-CreN-IRES2 using the primers listed in Table S1.

### Quail embryo EC culture

Fertilised quail eggs were incubated horizontally at 37.5°C in a humidified egg incubator until the desired developmental stages were reached. A hole was made at the blunt end of the eggs, and 2 mL albumin was removed using a 5 mL syringe (Nipro) fitted with an 18-gauge needle (Becton Dickinson) to lower the liquid level in the egg, ensuring the embryo was not disturbed during windowing. A small window (∼15 mm circle) was made along the long axis of the eggshell to facilitate visualisation and staging. Embryos were staged according to the H&H stage chart modified specifically for quail embryos (Ainsworth et al., 2010, Hamburger and Hamilton, 1951). Staged embryos were cooled for approximately 1 hour at room temperature (RT) or for 10-15 min at 4°C before EC culture. A cut was made at the pointed end of the egg, and the yolk was then decanted into the lid of a 90 mm Petri dish, with the embryo located on top of the yolk. The remaining albumin was gently removed using a 3 mL transfer pipette, and excess albumin from the surface of the yolk was gently blotted using Kimwipes. A piece of 26 mm x 26 mm square Whatman filter paper with an 11 mm x 3 mm hole in the centre was placed on top of the embryo, and all edges were gently pressed using fine forceps (Fine Science Tools) to ensure that the vitelline membrane adhered to the paper. All four edges of the filter paper were cut using fine scissors (Fine Science Tools), and the embryo was gently lifted away from the yolk at a 45° angle using fine forceps. The embryo was then placed ventral side up onto a 35 mm 1-well dish pre-coated with 1 mL albumin-agarose mixture consisting of 50% volume/volume quail albumin and 0.3% weight/volume bacto-agar (Becton Dickinson). The embryo was rinsed with pre-warmed 1xHanks’ Balanced Salt Solution (HBSS) using a 1 mL transfer pipette to remove excess yolk and transferred to a clean, coated 35 mm 1-well dish. Before further use, the 1-well dish containing the embryo was kept in a humidified environmental chamber at RT.

### Quail embryo *ex ovo* electroporation

Embryos were first incubated to the desired stage and cultured using the EC culture method described above. The negative electrode (Nepagene, CUY701P2E) is a 90 mm glass Petri dish with a platinum foil sheet (2 mm x 2 mm) embedded in the centre. The positive electrode (Nepagene, CUY701P2L) is a hand-held stem with a flat platinum paddle (2 mm x 2 mm) at the tip. Both electrodes were connected to the electroporator (Nepagene, NEPA21) using platinum wires. Before electroporation, the glass Petri dish was filled with approximately 32-35 mL of 1xHBSS, and the depth of the solution was around 5-7 mm. The injection needle was prepared from a glass micropipette (Drummond) that was pre-pulled using a micropipette puller (Sutter Instrument, model P-97). The injection needle was inserted into the capillary holder of a CellTram Oil microinjector (Eppendorf), and the tip of the needle was trimmed using fine scissors. The injection mix, containing the plasmid construct, sucrose solution and FastGreen FCF indicator solution (Merck) was aspirated into the injection needle. The final concentrations of vector DNA, sucrose and indicator solution were 4.2-5.8 µg/µL, 6% and 0.05%, respectively. The cultured embryo was placed at the centre of the platinum foil sheet electrode with the ventral side facing upwards. The tip of the needle was inserted into the embryonic tissue, and approximately 0.1 µL of the injection mix was injected beneath the desired region but above the vitelline membrane. A series of square electrical pulses (five pulses, 5.2 volts per pulse, with a 50 msec pulse length and 200 msec resting time) was applied. The embryo was then lifted and gently transferred ventral side up onto a clean, coated 35 mm 1-well dish. The electroporated embryos were returned to the 37.5°C humidified egg incubator overnight.

### Quail embryo *in ovo* electroporation

Quail embryos were electroporated *in ovo* at 5 to 6-somite stage (ss) using a previously described technique (McKeown et al., 2005), with some minor modifications. The eggs were windowed as described above, and approximately 0.2 mL of calligraphy ink (diluted 1:25 in 1xHBSS) was injected beneath the embryos using a 5 mL syringe fitted with a 27-gauge needle to facilitate visualisation. The injection mix was prepared as described above, and approximately 0.1 µL was injected into the neural tube lumen from both anterior and posterior neuropores to fill up the entire lumen. One to two drops of 1xHBSS were applied to the surface of the embryo to increase the electrical conductivity. The electrodes are approximately 40 mm platinum rods housed in a pen-like structure. Both platinum rods were bent approximately 6 mm from the exposed end at 90°, forming an L-shaped end. The platinum electrodes were placed on both side of the embryo, approximately 4-5 mm apart, with the negative electrode on the left side. Embryos were electroporated with three pulses of 25 volts, with a 50 msec pulse length and 200 msec resting time, using a custom-made electroporator (by Tatjana Sauka-Spengler). The window of electroporated embryos were sealed with Parafilm and returned to the 37.5°C humidified egg incubator overnight.

### Cre mRNA LNPs injection

Cre mRNA LNPs were a generous gift from Johnston lab (Chen et al., 2025), and were formulated in 25 mM Tris buffer with approximately 8.8% sucrose. The eggs were windowed and visualised with calligraphy ink injection as described above. Cre mRNA LNPs were mixed with Fast Green FCF indicator solution, with final concentrations of 260 ng and 0.05% for LNPs and indicator solution, respectively, and aspirated into the injection needle. Approximately 0.1 µL of injection mix was injected into the interstitial space between the posterior primitive streak and vitelline membrane (mesodermal cell labelling), the neural tube lumen (neuroepithelial and neural crest cell labelling) or the yolk sac vasculature (endothelial and circulating cell labelling). The windows of embryos were sealed with Parafilm and returned to the 37.5°C humidified egg incubator overnight.

## Whole-mount quail embryo immunofluorescent staining

Embryos were fixed at room temperature for 30 min in 4% paraformaldehyde (PFA, ProSciTech) and subsequently washed with phosphate-buffer saline (PBS) at RT three times, 5 min each time. Excess extra-embryonic tissues were trimmed before immunostaining. To enhance antibody penetration, the fixed embryos were permeabilised in PBS containing 0.5% Triton X-100 (PBTX, Sigma Aldrich) at RT three times, 30 min each time. The embryos were then blocked for 3 hours at RT in blocking buffer consisting of 0.5% PBTX, 1% bovine serum albumin (BSA, Sigma Aldrich) and 0.02% Sodium Dodecyl Sulfate (SDS, Sigma Aldrich) in PBS. After blocking, embryos were incubated with primary antibodies diluted in dilution buffer (0.5% PBTX, 0.2% BSA and 0.02% SDS in PBS) overnight at 4°C. Primary antibodies used were: Brachyury (In Vitro Technologies, AF2085-SP, 1:40), Cre recombinase (NEB, 15036T, 1:100), Fibronectin (DSHB, B3/D6-s, 1:5), Sox2 (Abcam, AB97959, 1:100), Sox10 (United Bioresearch, 66786-1-Ig, 1:1000). After primary antibody incubation, unbound antibodies were washed away with 0.5% PBTX at RT six times, 10 min each time. Embryos were then incubated with corresponding secondary antibodies (Alexa Fluor 647, Invitrogen, 1:1000) overnight at 4°C, and unbound secondary antibodies were washed away with 0.5% PBTX at RT six times, 10 min each time. To facilitate the imaging of deeper tissues, embryos were cleared sequentially in 30%, 60% and 90% glycerol in PBS for 10 min at each concentration and mounted on microscope slides with coverslips using 90% glycerol as the mounting medium. To visualise morphology without immunostaining, embryos were stained with Phalloidin-iFluor 647, (Abcam, AB176759, 1:1000) diluted in PBS overnight at 4°C, and unbound molecules were washed away with PBS three times, 10 min each time.

### Opto-Cre photoactivation

Electroporated embryos were transferred dorsal side down to a 6-well glass bottom plate (Cellvis, P06-1.5H-N) pre-coated with 250 µL albumin-agarose mixture. Opto-Cre photoactivation was performed with a Zeiss LSM710 confocal microscope equipped with LD C-Apo, 40x/1.1 Water objective (WD = 0.62 mm). A region of interest (ROI; ∼ 85 µm x 22 µm) was drawn on the epidermis using the rectangle tool in Zen software and exposed to 10 min irradiation with a 30% 488 nm laser. The embryos were returned to the 37.5°C humidified egg incubator overnight and imaged again to assess recombination by detecting nuclear EGFP-positive cells.

### Confocal imaging and vibratome sectioning

Whole-mount immunostained embryos were imaged using a Zeiss LSM710 confocal microscope equipped with GaAsP detectors using either PlnApo, 20x/0.8 DIC II objective (WD = 0.55 mm) or LD C-Apo 40x/1.1 Water objective. To visualise both dorsal root ganglion (DRG) and sympathetic ganglion (SG), embryos were subjected to vibratome sectioning. After whole-mount imaging, embryos were removed from the microscope slide and subjected sequentially to 90%, 60%, 30% glycerol and PBS for 10 min at each condition. Embryos were then embedded in Tissue-Tek Cryomolds (ProSciTech, Biopsy) containing 5% agarose and 15% sucrose in PBS. Embedded embryos were sectioned using a Leica VT1000 S vibrating blade microtome at a thickness of 80 µm. The sectioned slices were subjected to clearing as described above before mounting on the microscope slides.

### Use of artificial intelligence tools

ChatGPT (OpenAI) was used to improve the grammar, clarity and readability of the manuscript. The authors reviewed and approved all AI-assisted edits and take full responsibility for the content of the manuscript.

## SUPPLEMENTARY DATA

**Fig S1.**
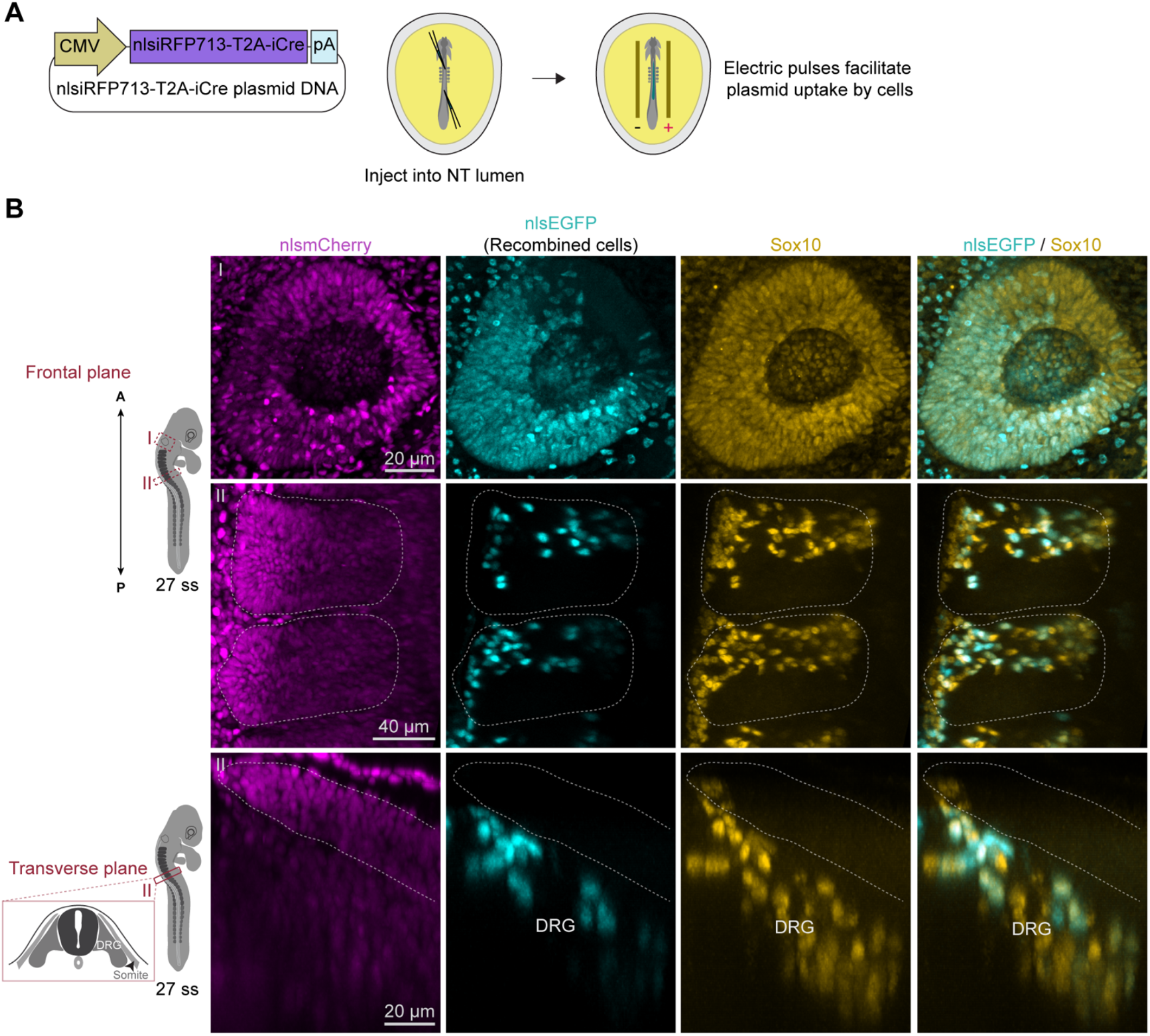
Fig S1. Cre-mediated recombination occurs in different tissues. (A) Plasmid vectors used for *in ovo* electroporation. (B) Following *in ovo* electroporation, Cre-mediated recombination activates nuclear EGFP expression in the otic vesicle (I) and the dorsal root ganglion (DRG). Sox10 immunostaining identifies a subset of the recombined nuclear EGFP-positive cells as neural crest cells (gold). Dotted lines indicate somites.

**Fig S2.**
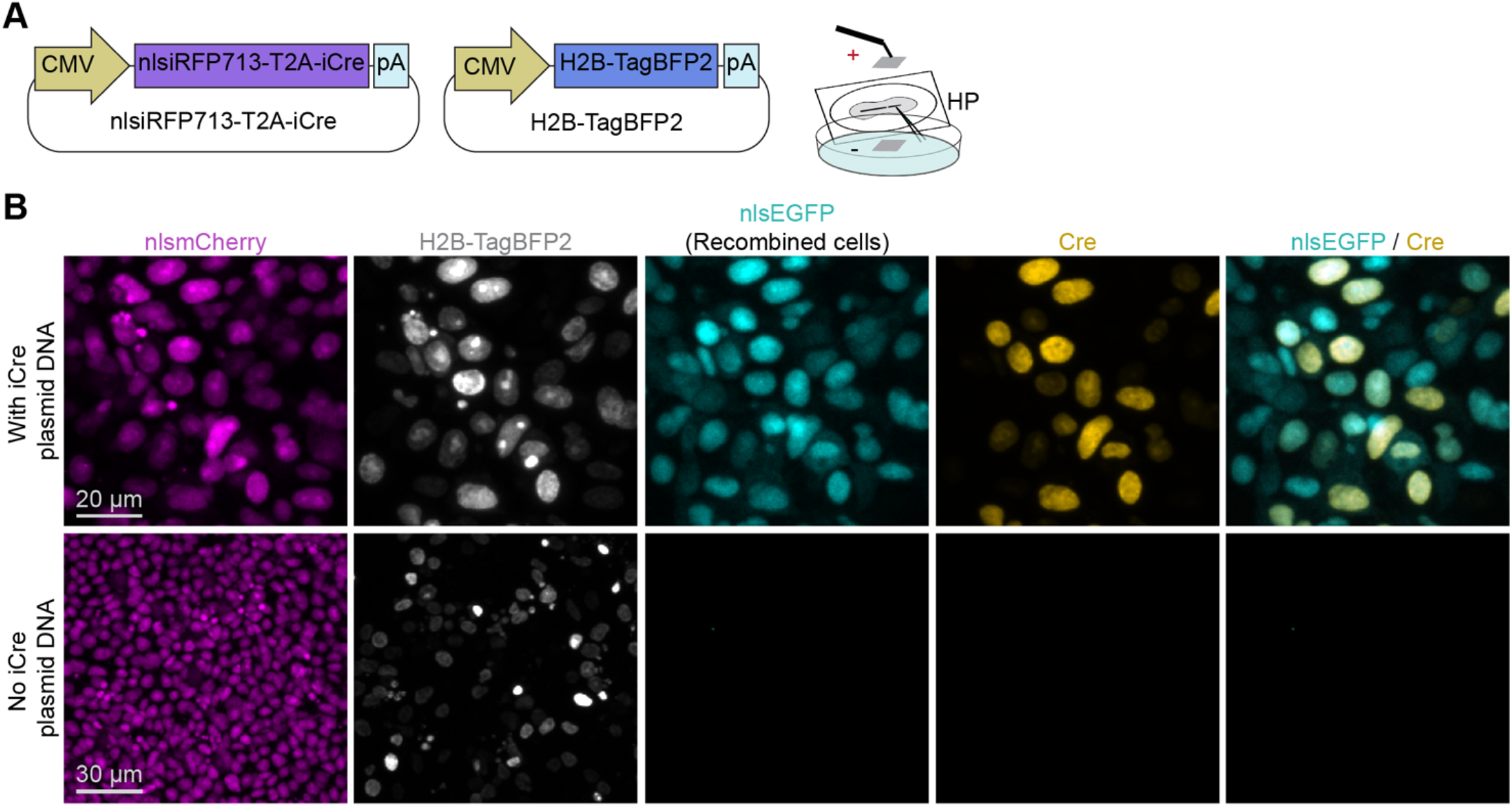
Validation of Cre-mediated recombination. (A) Plasmid vectors used for *ex ovo* electroporation. (B) In the presence of iCre plasmid, most recombined nuclear EGFP-positive cells express Cre (top panel), whereas in the absence of iCre plasmid, no Cre expression or recombination is observed (bottom panel).

**Fig S3.**
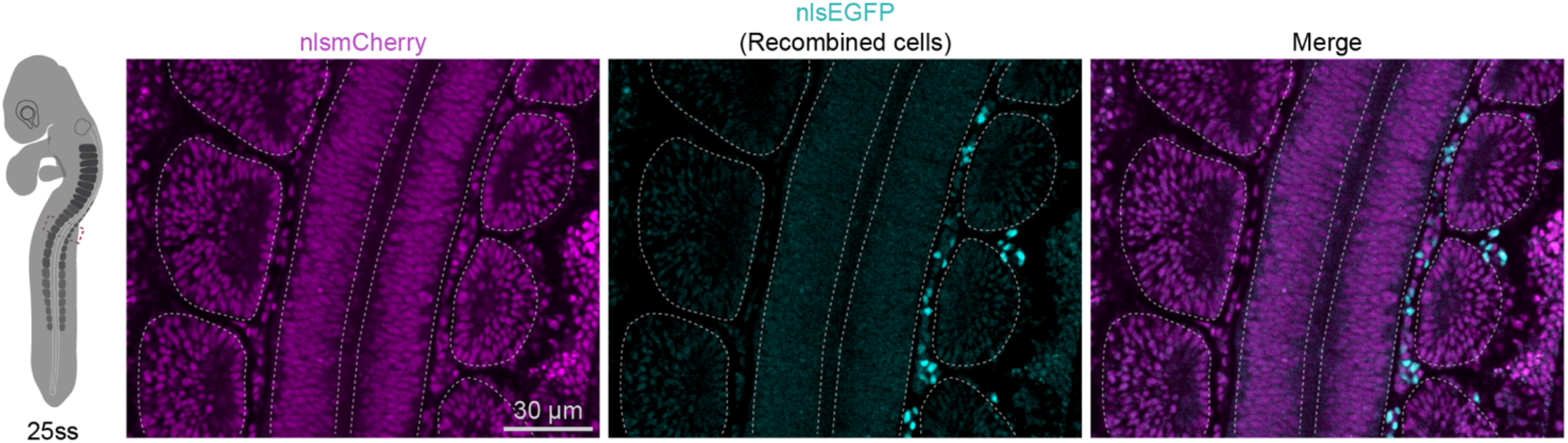
Unilateral *in ovo* electroporation. Embryos were electroporated unilaterally with the *FoxD3* NC2-iCre plasmid at 6ss and fixed after overnight incubation. A representative embryo at 25ss shows that nuclear EGFP-positive cells are predominantly detected on the right side, the side of electroporation.

**Fig S4.**
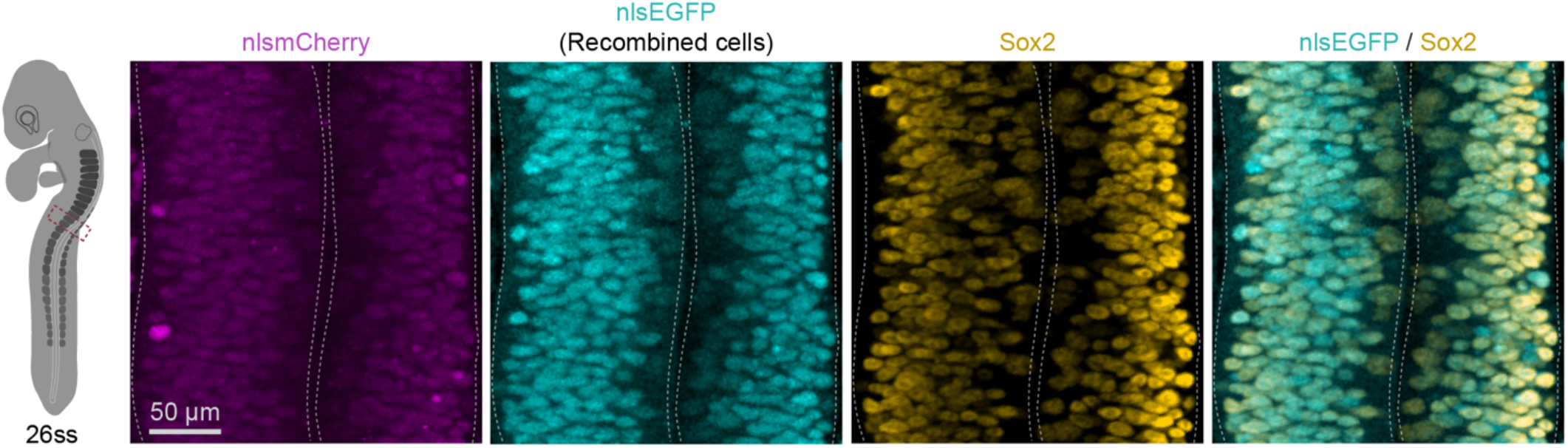
Cre mRNA LNP-mediated recombination in the neuroepithelium. Embryos were injected with Cre mRNA LNPs into the neural tube lumen at 5ss and fixed after overnight incubation. A representative embryo at 26ss shows that most recombined nuclear EGFP-positive cells are Sox2-positive, confirming their neuroepithelial identity.

**Fig S5.**
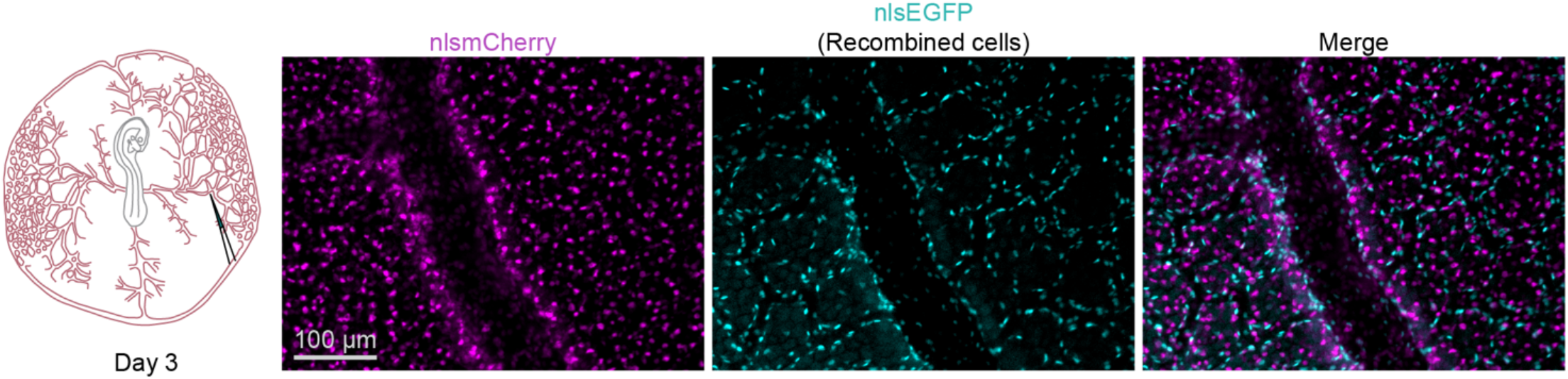
Cre mRNA LNP-mediated recombination in the yolk sac vasculature. Cre mRNA LNPs were injected into the yolk sac vasculature at day 3 of incubation. Following overnight incubation, Cre mRNA LNP-mediated recombination resulted in nuclear EGFP expression in endothelial cells and circulating cells.

**Table S1.**
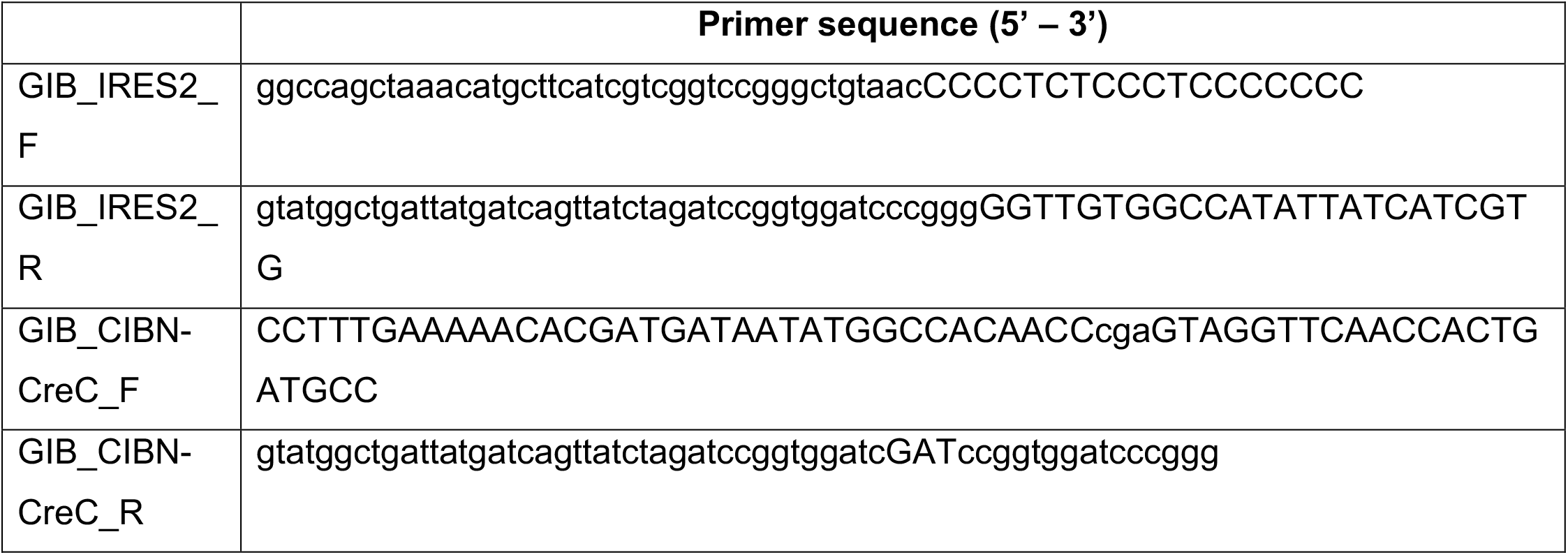
Primer sequences used to generate the opto-Cre plasmid construct.

## ACKNOWLEDGEMENTS

The authors acknowledge the IMB Advanced Microscopy Platform for access to specialised optical imaging instrumentation, associated computational analysis resources, and expert staff support.

## DATA AND RESOURCE AVAILABILITY

All relevant data and details of resources can be found within the article and its supplementary information.

## REFERENCES

Ainsworth, S. J., Stanley, R. L. & Evans, D. J. 2010. Developmental stages of the Japanese quail. J Anat, 216, 3–15.

Alvarez, Y. D., Van Der Spuy, M., Wang, J. X., Noordstra, I., Tan, S. Z., Carroll, M., Yap, A. S., Serralbo, O. & White, M. D. 2024. A Lifeact-EGFP quail for studying actin dynamics in vivo. Journal of Cell Biology, 223.

Azambuja, A. P. & Simões-Costa, M. 2021. The connectome of neural crest enhancers reveals regulatory features of signaling systems. Dev Cell, 56, 1268–1282.e6.

Barzilai-Tutsch, H., Morin, V., Toulouse, G., Chernyavskiy, O., Firth, S., Marcelle, C. & Serralbo, O. 2022. Transgenic quails reveal dynamic TCF/β-catenin signaling during avian embryonic development. eLife, 11, e72098.

Betancur, P., Bronner-Fraser, M. & Sauka-Spengler, T. 2010a. Assembling Neural Crest Regulatory Circuits into a Gene Regulatory Network. Annual Review of Cell and Developmental Biology, 26, 581–603.

Betancur, P., Bronner-Fraser, M. & Sauka-Spengler, T. 2010b. Genomic code for Sox10 activation reveals a key regulatory enhancer for cranial neural crest. Proc Natl Acad Sci U S A, 107, 3570–5.

Caldarelli, P., Chamolly, A., Villedieu, A., Alegria-Prévot, O., Phan, C., Gros, J. & Corson, F. 2024. Self-organized tissue mechanics underlie embryonic regulation. Nature, 633, 887–894.

Chen, M. Z., Yuen, D., Mcleod, V. M., Yong, K. W., Smyth, C. H., Herling, B. R., Payne, T. J., Fabb, S. A., Belousoff, M. J., Algarni, A., Sexton, P. M., Porter, C. J. H., Pouton, C. W. & Johnston, A. P. R. 2025. A versatile antibody capture system drives specific in vivo delivery of mRNA-loaded lipid nanoparticles. Nature Nanotechnology, 20, 1273–1284.

Chinnaiya, K., Burbridge, S., Jones, A., Kim, D. W., Place, E., Manning, E., Groves, I., Sun, C., Towers, M., Blackshaw, S. & Placzek, M. 2023. A neuroepithelial wave of BMP signalling drives anteroposterior specification of the tuberal hypothalamus. Elife, 12.

Gandhi, S., Li, Y., Tang, W., Christensen, J. B., Urrutia, H. A., Vieceli, F. M., Piacentino, M. L. & Bronner, M. E. 2021. A single-plasmid approach for genome editing coupled with long-term lineage analysis in chick embryos. Development, 148.

Gandhi, S., Rajan, A. R. D., Urrutia, H., Wilson, J. S., Ling, I. T. C., Williams, R., Sauka-Spengler, T. & Bronner, M. E. 2026. Regulatory logic underlying neural crest contributions to the head versus the heart. Proc Natl Acad Sci U S A, 123, e2512031123.

Graham, A. 2003. The neural crest. Current Biology, 13, R381-R384.

Hadas, Y., Etlin, A., Falk, H., Avraham, O., Kobiler, O., Panet, A., Lev-Tov, A. & Klar, A. 2014. A ‘tool box’ for deciphering neuronal circuits in the developing chick spinal cord. Nucleic Acids Research, 42, e148–e148.

Hamburger, V. & Hamilton, H. L. 1951. A series of normal stages in the development of the chick embryo. Journal of Morphology, 88, 49–92.

Harn, H. I., Jiang, T. X., Huang, C. H., Juan, W. T., Liu, T. Y., Chuang, T. C., Liao, W. C., Wang, Y., Li, J., Weijer, C. J., Wu, P., Guo, C. L. & Chuong, C. M. 2026. Novel tissue mechanics-guided cellular flows drive the formation of feather follicles. Embo j, 45, 3926–3953.

Hartwich, H., Satheesh, S. V. & Nothwang, H. G. 2012. A pink mouse reports the switch from red to green fluorescence upon Cre-mediated recombination. BMC Res Notes, 5, 296.

Hitoshi, N., Ken-Ichi, Y. & Jun-Ichi, M. 1991. Efficient selection for high-expression transfectants with a novel eukaryotic vector. Gene, 108, 193–199.

Huss, D., Benazeraf, B., Wallingford, A., Filla, M., Yang, J., Fraser, S. E. & Lansford, R. 2015. A transgenic quail model that enables dynamic imaging of amniote embryogenesis. Development, 142, 2850–9.

Jacobs-Li, J., Tang, W., Li, C. & Bronner, M. E. 2023. Single-cell profiling coupled with lineage analysis reveals vagal and sacral neural crest contributions to the developing enteric nervous system. eLife, 12, e79156.

Kohl, A., Hadas, Y., Klar, A. & Sela-Donenfeld, D. 2013. Electroporation of the hindbrain to trace axonal trajectories and synaptic targets in the chick embryo. J Vis Exp, e50136.

Le Douarin, N. 1973. A biological cell labeling technique and its use in experimental embryology. Developmental Biology, 30, 217–222.

Loulier, K., Barry, R., Mahou, P., Le franc, Y., Supatto, W., Matho Katherine s., Ieng, S., Fouquet, S., Dupin, E., Benosman, R., Chédotal, A., Beaurepaire, E., Morin, X. & Livet, J. 2014. Multiplex Cell and Lineage Tracking with Combinatorial Labels. Neuron, 81, 505–520.

Mckeown, S. J., Lee, V. M., Bronner-Fraser, M., Newgreen, D. F. & Farlie, P. G. 2005. Sox10 overexpression induces neural crest-like cells from all dorsoventral levels of the neural tube but inhibits differentiation. Developmental Dynamics, 233, 430–444.

Ming, Z., Liu, F., Moran, H. R., Lalonde, R. L., Adams, M., Restrepo, N. K., Joshi, P., Ekker, S. C., Clark, K. J., Friedberg, I., Sumanas, S., Yin, C., Mosimann, C., Essner, J. J. & Mcgrail, M. 2026. Lineage labeling with zebrafish hand2 Cre and CreERT2 recombinase CRISPR knock-ins. Developmental Dynamics, 255, 86–105.

Morris, K. M., Hindle, M. M., Boitard, S., Burt, D. W., Danner, A. F., Eory, L., Forrest, H. L., Gourichon, D., Gros, J., Hillier, L. W., Jaffredo, T., Khoury, H., Lansford, R., Leterrier, C., Loudon, A., Mason, A. S., Meddle, S. L., Minvielle, F., Minx, P., Pitel, F., Seiler, J. P., Shimmura, T., Tomlinson, C., Vignal, A., Webster, R. G., Yoshimura, T., Warren, W. C. & Smith, J. 2020. The quail genome: insights into social behaviour, seasonal biology and infectious disease response. BMC Biol, 18, 14.

Murko, C. & Bronner, M. E. 2017. Tissue specific regulation of the chick Sox10E1 enhancer by different Sox family members. Dev Biol, 422, 47–57.

Nitzan, E., Pfaltzgraff, E. R., Labosky, P. A. & Kalcheim, C. 2013. Neural crest and Schwann cell progenitor-derived melanocytes are two spatially segregated populations similarly regulated by Foxd3. Proc Natl Acad Sci U S A, 110, 12709–14.

Oh, J. D. H., Freem, L., Saunders, D. D. Z., Mcteir, L., Gilhooley, H., Jackson, M., Glover, J. D., Smith, J., Schoenebeck, J. J., Lettice, L. A., Sang, H. M. & Davey, M. G. 2024. Insights into digit evolution from a fate map study of the forearm using Chameleon, a new transgenic chicken line. Development, 151.

Pfann, M., Ben-Tal Cohen, E., Sela-Donenfeld, D. & Cinnamon, Y. 2025. Application of the Magnet-Cre optogenetic system in the chicken model. Developmental Biology, 523, 68–81.

Ribatti, D. 2019. Nicole Le Douarin and the use of quail-chick chimeras to study the developmental fate of neural crest and hematopoietic cells. Mechanisms of Development, 158, 103557.

Schick, E., Mccaffery, S. D., Keblish, E. E., Thakurdin, C. & Emerson, M. M. 2019. Lineage tracing analysis of cone photoreceptor associated cis-regulatory elements in the developing chicken retina. Sci Rep, 9, 9358.

Serralbo, O., Salgado, D., Véron, N., Cooper, C., Dejardin, M.-J., Doran, T., Gros, J. & Marcelle, C. 2020. Transgenesis and web resources in quail. eLife, 9, e56312.

Shaner, N. C., Campbell, R. E., Steinbach, P. A., Giepmans, B. N. G., Palmer, A. E. & Tsien, R. Y. 2004. Improved monomeric red, orange and yellow fluorescent proteins derived from Discosoma sp. red fluorescent protein. Nature Biotechnology, 22, 1567–1572.

Simões-Costa, M. S., Mckeown, S. J., Tan-Cabugao, J., Sauka-Spengler, T. & Bronner, M. E. 2012. Dynamic and Differential Regulation of Stem Cell Factor FoxD3 in the Neural Crest Is Encrypted in the Genome. PLOS Genetics, 8, e1003142.

Takahashi, M., Isagawa, T., Sato, T., Takeda, N. & Kawakami, K. 2024. Lineage tracing using Wnt2b-2A-CreERT2 knock-in mice reveals the contributions of Wnt2b-expressing cells to novel subpopulations of mesothelial/epicardial cell lineages during mouse development. Genes to Cells, 29, 854–875.

Tombácz, I., Laczkó, D., Shahnawaz, H., Muramatsu, H., Natesan, A., Yadegari, A., Papp, T. E., Alameh, M. G., Shuvaev, V., Mui, B. L., Tam, Y. K., Muzykantov, V., Pardi, N., Weissman, D. & Parhiz, H. 2021. Highly efficient CD4+ T cell targeting and genetic recombination using engineered CD4+ cell-homing mRNA-LNPs. Mol Ther, 29, 3293–3304.

Tyack, S. G., Jenkins, K. A., O’neil, T. E., Wise, T. G., Morris, K. R., Bruce, M. P., Mcleod, S., Wade, A. J., Mckay, J., Moore, R. J., Schat, K. A., Lowenthal, J. W. & Doran, T. J. 2013. A new method for producing transgenic birds via direct in vivo transfection of primordial germ cells. Transgenic Research, 22, 1257–1264.

Villedieu, A., Alegria-Prévot, O., Phan, C., Ieda, Y., Corson, F. & Gros, J. 2025. Live imaging and functional characterization of the avian hypoblast redefine the mechanisms of primitive streak induction. Nature Communications, 16, 11616.

Wang, J. X., Alvarez, Y. D., Tan, S. Z., Ranie, S. N., Stehbens, S. J. & White, M. D. 2026. Quantitative live imaging reveals PRICKLE1 controls junctional neural tube morphogenesis independent of Planar Cell Polarity. Nature Communications, 17, 3654.

Williams, R. M., Candido-Ferreira, I., Repapi, E., Gavriouchkina, D., Senanayake, U., Ling, I. T. C., Telenius, J., Taylor, S., Hughes, J. & Sauka-Spengler, T. 2019. Reconstruction of the Global Neural Crest Gene Regulatory Network In Vivo. Dev Cell, 51, 255–276.e7.

Yoshinari, N., Ando, K., Kudo, A., Kinoshita, M. & Kawakami, A. 2012. Colored medaka and zebrafish: Transgenics with ubiquitous and strong transgene expression driven by the medaka β-actin promoter. Development, Growth & Differentiation, 54, 818–828.

